# Omicron-induced interferon signalling prevents influenza A virus infection

**DOI:** 10.1101/2022.09.06.506799

**Authors:** Denisa Bojkova, Marco Bechtel, Tamara Rothenburger, Joshua D. Kandler, Lauren Hayes, Ruth Olmer, Martin Ulrich, Danny Jonigk, Sandra Ciesek, Mark N. Wass, Martin Michaelis, Jindrich Cinatl

## Abstract

Recent findings in permanent cell lines suggested that SARS-CoV-2 Omicron BA.1 induces a stronger interferon response than Delta. Here, we show that BA.1 and BA.5 but not Delta induce an antiviral state in air-liquid interface (ALI) cultures of primary human bronchial epithelial (HBE) cells and primary human monocytes. Both Omicron subvariants caused the production of biologically active type I (α/β) and III (λ) interferons and protected cells from super-infection with influenza A viruses. Notably, abortive Omicron infection of monocytes was sufficient to protect monocytes from influenza A virus infection. Interestingly, while influenza-like illnesses surged during the Delta wave in England, their spread rapidly declined upon the emergence of Omicron. Mechanistically, Omicron-induced interferon signalling was mediated via double-stranded RNA recognition by MDA5, as MDA5 knock-out prevented it. The JAK/ STAT inhibitor baricitinib inhibited the Omicron-mediated antiviral response, suggesting it is caused by MDA5-mediated interferon production, which activates interferon receptors that then trigger JAK/ STAT signalling. In conclusion, our study 1) demonstrates that only Omicron but not Delta induces a substantial interferon response in physiologically relevant models, 2) shows that Omicron infection protects cells from influenza A virus super-infection, and 3) indicates that BA.1 and BA.5 induce comparable antiviral states.

## Introduction

SARS-CoV-2, the coronavirus that causes COVID-19, has caused the worst pandemic since the Spanish Flu in 1918-1920 [Bastard et al., 2022]. Virus-induced interferon signalling has been shown to be critically involved in determining COVID-19 severity [Bastard et al., 2022; Chiale et al., 2022]. Individuals with defects in their interferon response are predisposed to life-threatening COVID-19 [Bastard et al., 2022; Chiale et al., 2022], and a particularly pronounced interferon-related innate immune response is anticipated to contribute to the lower COVID-19 severity observed in children [Borrelli et al., 2021].

Recent findings suggested that SARS-CoV-2 Omicron BA.1 displays a lower interferon antagonism than Delta [Bojkova et al., 2022; Bojkova et al., 2022a]. BA.1 and Delta viruses showed a similar replication pattern in interferon-deficient Vero cells, but BA.1 replication was attenuated relative to Delta in interferon competent Calu-3, Caco-2, and Caco-2-F03 (a highly SARS-CoV-2-susceptible Caco-2 subline [Bojkova et al., 2022b]) cells [Bojkova et al., 2022; Hu et al., 2022; Shuai et al., 2022]. Moreover, BA.1 induced a more pronounced interferon response than Delta [Bojkova et al., 2022; Bojkova et al., 2022a].

Here, we investigated Delta, BA.1, and BA.5 replication in air-liquid interface (ALI) cultures of primary human bronchial epithelial (HBE) cells and primary human monocytes. To examine whether SARS-CoV-2-induced interferon induction results in a biologically relevant antiviral state, we further determined the impact of Delta, BA.1, and BA.5 infection on influenza A virus replication in ALI HBE cultures (H1N1) and monocytes (H1N1, H5N1), as interferon signalling is considered to be a major influenza A virus restriction factor [McKellar et al., 2021].

## Results

### Infection kinetics and interferon response induction by Omicron BA.1 and Delta in air-liquid-interface (ALI) cultures of primary human bronchial epithelial (HBE) cells

BA.1 displayed faster replication kinetics than Delta in ALI HBE cultures (Figure 1), as indicated by high SARS-CoV-2 nucleoprotein (NP), genomic RNA levels, and caspase 3/7 activity (which reflects SARS-CoV-2 replication independently of whether the virus causes cytotoxicity resulting in a cytopathogenic effect in a cell culture model [Bojkova et al., 2022b]) (Figure 1B-D). However, the replication of both SARS-CoV-2 variants resulted in comparable peak NP and genomic RNA levels (Figure 1B-D). While the BA.1 levels declined after a peak (at 24h post infection for NP and 72h for genomic RNA), Delta levels continued to increase until 120h post infection (Figure 1B-D). These findings are in accordance with previous findings showing that BA.1 replicates faster than other variants in bronchial cells [Hui et al., 2022]. Independently of the replication kinetics BA.1 and Delta caused similar reductions of the ALI HBE barrier integrity (Figure 1E).

**Figure 1.**
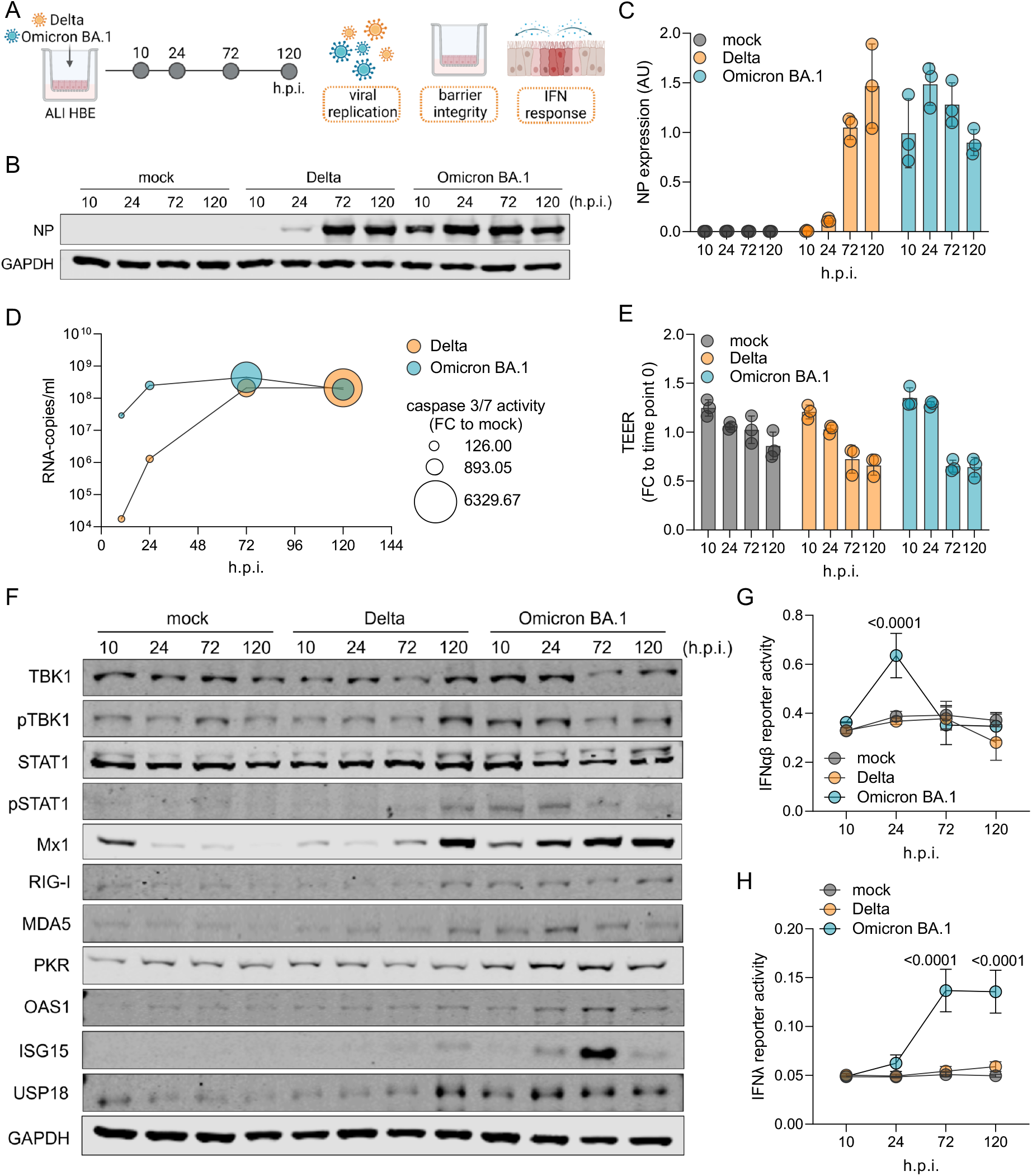
Infection kinetics and interferon response induction by Omicron BA.1 and Delta in air-liquid-interface (ALI) cultures of primary human bronchial epithelial (HBE) cells. (A) Schematic depiction of the experimental set-up. (B) Immunoblot of cellular SARS-CoV-2 nucleoprotein (NP) levels in BA.1- and Delta (MOI 1)-infected ALI HBE cultures at different time points post infection. (C) Quantification of NP levels (mean ± SD) by ImageJ. (D) SARS-CoV-2 genomic RNA copy numbers (Y axis) and caspase 3/7 activity (bubble size) determined in the apical medium of BA.1- and Delta (MOI 1)-infected at different time points post infection. (E) Evaluation of barrier integrity by measurement of TEER in BA.1- and Delta (MOI 1)-infected ALI HBE cultures at different time points post infection. Bars represents mean ± SD of three biological replicates. (F) Immunoblot indicating cellular levels of proteins involved in interferon signalling in BA.1- and Delta (MOI 1)-infected ALI HBE cultures at different time points post infection. (G,H) Interferon (IFN)-α/β (G) and IFNλ (H) responses induced in HEK reporter cell lines by apical washes from BA.1- and Delta (MOI 1)-infected ALI HBE cultures collected at different time points post infection. P values were determined by one-way ANOVA with subsequent Tukey’s test.

Also in agreement with previous findings [Alfi et al., 2022; Bojkova et al., 2022; Bojkova et al., 2022a], BA.1 induced a stronger interferon response than Delta, as indicated by the abundance and phosphorylation levels of a range of proteins involved in interferon signalling (Figure 1F). Moreover, only BA.1 but not Delta induced the secretion of biologically active interferon-α/β (Figure 1G) and -λ (Figure 1H) by ALI HBE cultures, as demonstrated using HEK-reporter cell lines.

Interferon-α/β peaked at 24h post infection (Figure 1G), which was followed by a return to basal levels, whereas interferon-λ remained elevated until 120h post infection (Figure 1H). Short-term interferon type I (α/β) responses cause a protective antiviral response, while long-term interferon activity is associated with potentially deleterious inflammation [Forero et al., 2019; King & Sprent, 2021]. In contrast, sustained interferon type III (λ) responses inhibit respiratory virus replication at epithelial barriers in the respiratory tract and prevent excessive inflammation [Forero et al., 2019; Prokunina-Olsson et al., 2020; King & Sprent, 2021]. Hence, the interferon type I and III responses observed in BA.1-infected ALI HBE cultures add further evidence explaining why Omicron is less pathogenic than other SARS-CoV-2 variants like Delta [Wang et al., 2022].

### JAK/ STAT inhibition suppresses BA.1-induced interferon signalling and increases BA.1 replication in air-liquid-interface (ALI) human bronchial epithelial (HBE) cell cultures

Interferon signalling can be induced in a STAT1-dependent and -independent manner [Rani et al., 2010; Bastard et al., 2022; Chiale et al., 2022]. In hibition of JAK/ STAT signalling using the JAK inhibitor baricitinib significantly increased BA.1 replication in ALI HBE cultures (Figure 2), as indicated by genomic RNA copy numbers (Figure 2B) and cellular NP levels (Figure 2C). In contrast, Delta only displayed a non-significant trend towards higher genomic RNA copy numbers (Figure 2B) and cellular NP levels (Figure 2C) in the presence of baricitinib. Baricitinib did not exert significant effects on BA.1- and Delta-mediated caspase 3/7 activation (Figure 2D) and the ALI HBE barrier function (Figure 2E), although there was a non-significant trend towards enhanced caspase 3/7 activity in BA.1-infected ALI HBE cultures (Figure 2D).

**Figure 2.**
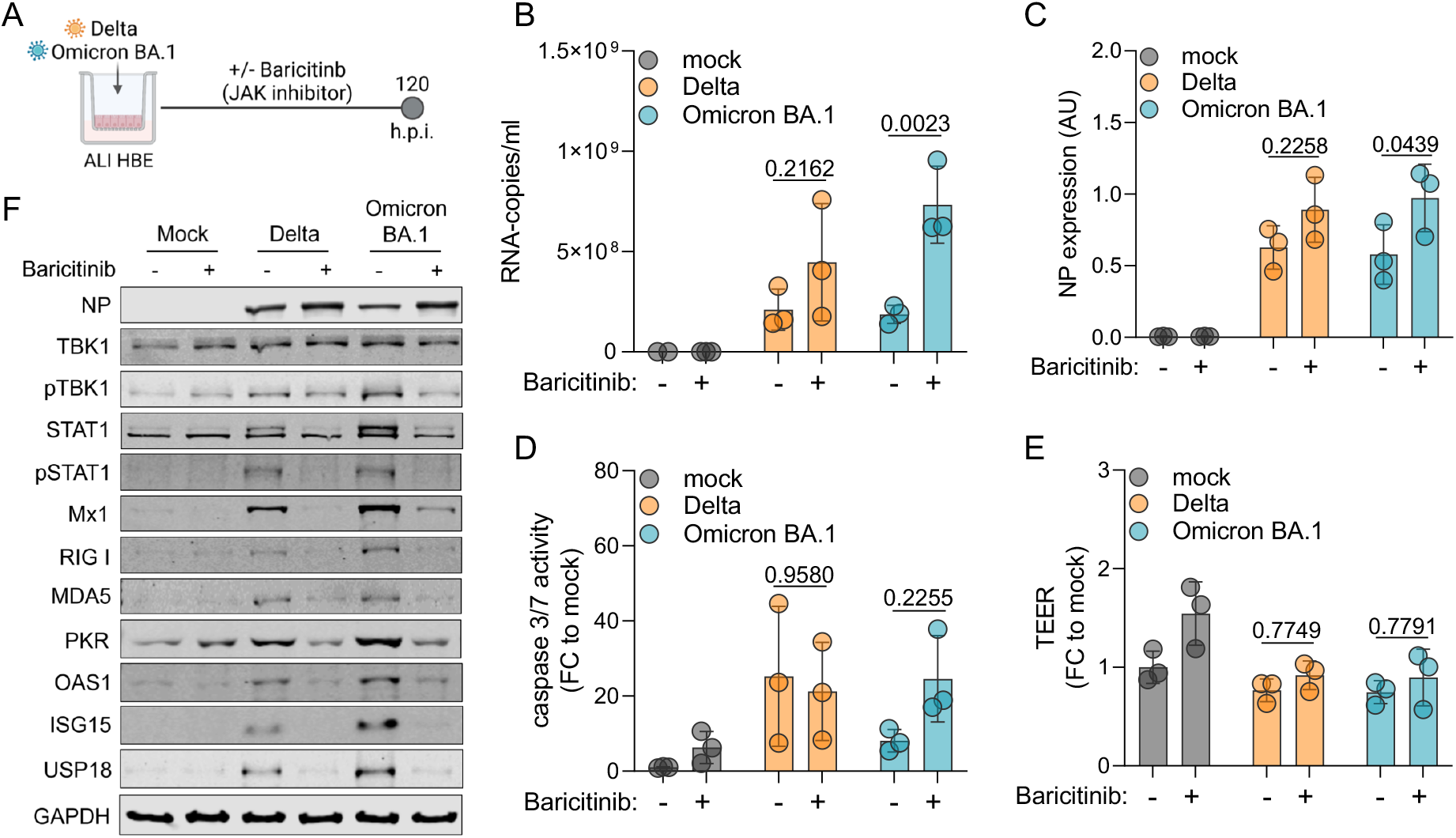
Inhibition of JAK/STAT signalling promotes Omicron BA.1 replication. (A) Schematic depiction of the experimental set-up. (B) SARS-CoV-2 genomic RNA copies in apical wash of air-liquid-interface (ALI) human bronchial epithelial (HBE) cultures 120 h post SARS-CoV-2 (MOI 1) infection in the absence or presence of the JAK inhibitor baricitinib (1μM). Bars represent mean ± SD of three biological replicates. (C) NP levels in SARS-CoV-2 (MOI 1)-infected ALI HBE cultures in the absence or presence of baricitinib (1μM) as determined by immunoblotting. Bars represent mean ± SD of three biological replicates. (D) Caspase 3/7 activation in apical washes of SARS-CoV-2 (MOI 1)-infected ALI HBE cultures in the absence or presence of baricitinib (1μM) 120h post infection. Bars display mean ± SD of three biological replicates. (E) Barrier integrity in SARS-CoV-2 (MOI 1)-infected ALI HBE cultures in the absence or presence of baricitinib (1μM) measured by TEER at 120h post infection. Mean ± SD of three biological replicates is presented. (F) Immunoblot indicating cellular levels of proteins involved in interferon signalling in SARS-CoV-2 (MOI 1)-infected ALI HBE cultures at 120h post infection in the presence or absence of baricitinib (1μM). All P values were calculated by Student’s t-test.

Western blot analysis confirmed that baricitinib not only increased BA.1 replication but also suppressed BA.1-induced interferon signalling (Figure 2F). Taken together, these findings indicate that the pronounced interferon response induced by BA.1 is mediated via STAT1 and that it attenuates BA.1 replication.

### BA.1-induced interferon signalling protects air-liquid-interface (ALI) human bronchial epithelial (HBE) cell cultures from H1N1 influenza A virus super-infection

Next, we here infected ALI HBE cultures with BA.1 or Delta (both MOI 1) for 48h prior to infection with H1N1 influenza A virus (MOI 2) (Figure 3A) to examine whether the BA.1-induced interferon response may induce an antiviral state that interferes with H1N1 replication. Controls confirmed that both BA.1 and Delta replication as well as BA.1- and Delta-induced interferon induction were comparable to the data presented in Figure 1 (Suppl. Figure 1).

**Figure 3.**
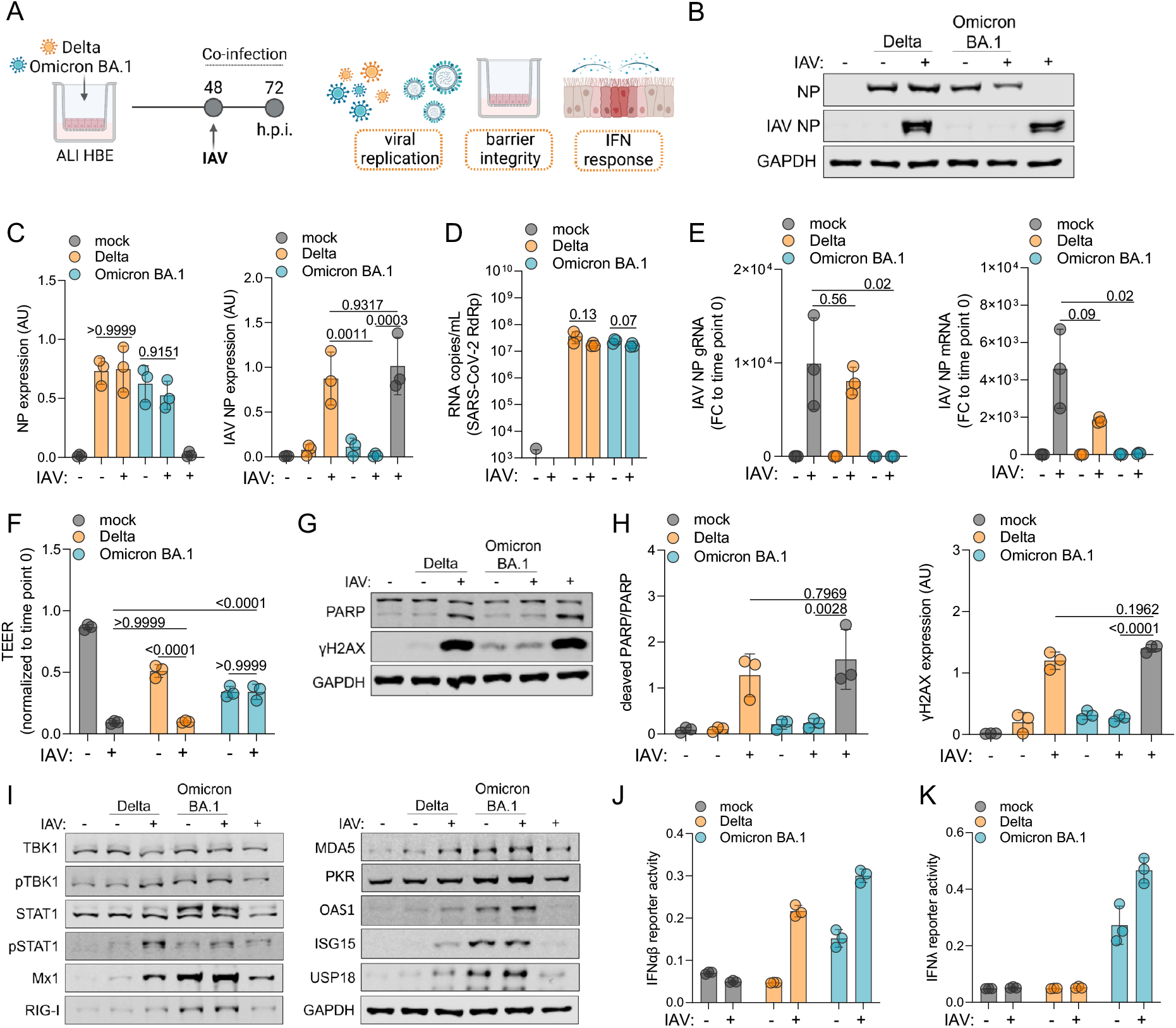
Omicron BA.1 but not Delta infection prevents H1N1 influenza A virus (IAV) replication. (A) Schematic of the experimental set-up. (B) Immunoblot of the SARS-CoV-2 nucleoprotein (NP) and H1N1 IAV NP levels 24 h post infection with H1N1 strain A/NewCaledonia/20/99 (MOI 2). (C) Quantification of immunoblots from (B). Bars represent mean ± SD of three biological replicates. P values were calculated by two-way ANOVA. (D) Genomic RNA copies of the SARS-CoV-2 RNA-dependent RNA polymerase (RdRp) in the apical wash of ALI HBE cultures 24 hours post infection with H1N1 (MOI 2). Values represent means ± SD of three biological replicates. P values were determined by Student’s t-test. (E) IAV NP genomic RNA (gRNA, left) or mRNA (right) levels 24h post infection with H1N1 (MOI 2). Bars display means ± SD of three biological replicates. P values were determined by Student’s t-test. (F) Barrier integrity measure by transepithelial electric resistance (TEER) in single- or co-infected ALI cultures. Bars display means ± SD of three biological replicates. P values were calculated by two-way ANOVA. (G) Immunoblot of PARP cleavage and γH2AX after single- or co-infection. (H) Quantification of the immunoblots from (G). Values represent the mean ± SD of three biological replicates. P values were calculated by two-way ANOVA. (I) Immunoblots displaying the levels of proteins involved in interferon signalling in single- and co-infected ALI HBE cultures. (J, K) Interferon (IFN)α/β (J) or IFNλ (K) signalling in HEK-reporter cell lines incubated with apical washes of single- and co-infected ALI HBE cultures 24 hours post infection with H1N1 (MOI 2).

Determination of H1N1 nucleoprotein (NP) levels indicated that only BA.1 but not Delta suppressed H1N1 infection in ALI HBE cultures (Figure 3B, Figure 3C). H1N1 super-infection did not significantly reduce SARS-CoV-2 levels in ALI HBE cultures (Figure 3D).

Genomic H1N1 NP RNA and H1N1 NP mRNA levels confirmed that only BA.1 caused a significant reduction of H1N1 replication (Figure 3E). Moreover, only BA.1 infection prevented H1N1-induced cytotoxicity as indicated by transepithelial electric resistance (TEER) measurement (Figure 3F). Influenza A virus replication is associated with the induction of apoptosis and DNA damage induction in host cells [Li et al., 2015; Ampomah & Lim, 2020], and PARP cleavage and γH2AX levels also confirmed that BA.1 infection suppressed H1N1-induced cytotoxicity (Figure 3G, Figure 3H).

The analysis of proteins involved in interferon signalling showed that also in the presence of H1N1 only BA.1 induced a pronounced interferon response (Figure 3I). In agreement, only BA.1-infected (but not of Delta-infected) ALI HBE cultures produced biologically active type I (α/ß) and III (λ) interferons, as demonstrated using HEK-reporter cell lines (Figure 3J, Figure 3K). Taken together, these findings indicate that only BA.1 but not Delta induces an interferon-mediated antiviral state in ALI-HBE cultures that protects them from H1N1 infection.

### Inhibition of JAK/ STAT signalling promotes H1N1 influenza A virus replication in BA.1-infected cells

Next, we investigated the effect of the JAK inhibitor on the BA.1-mediated suppression of H1N1 replication (Figure 4A). In agreement with the data presented in Figure 2 and Figure 3, baricitinib increased BA.1 replication as indicated by genomic RNA copies of the viral RNA-dependent RNA polymerase gene (Figure 4B). Moreover, baricitinib prevented the BA.1-mediated inhibition of H1N1 replication as indicated by H1N1 NP mRNA and genomic RNA levels (Figure 4C) and H1N1 NP protein levels (Figure 4D, Figure 4E).

**Figure 4.**
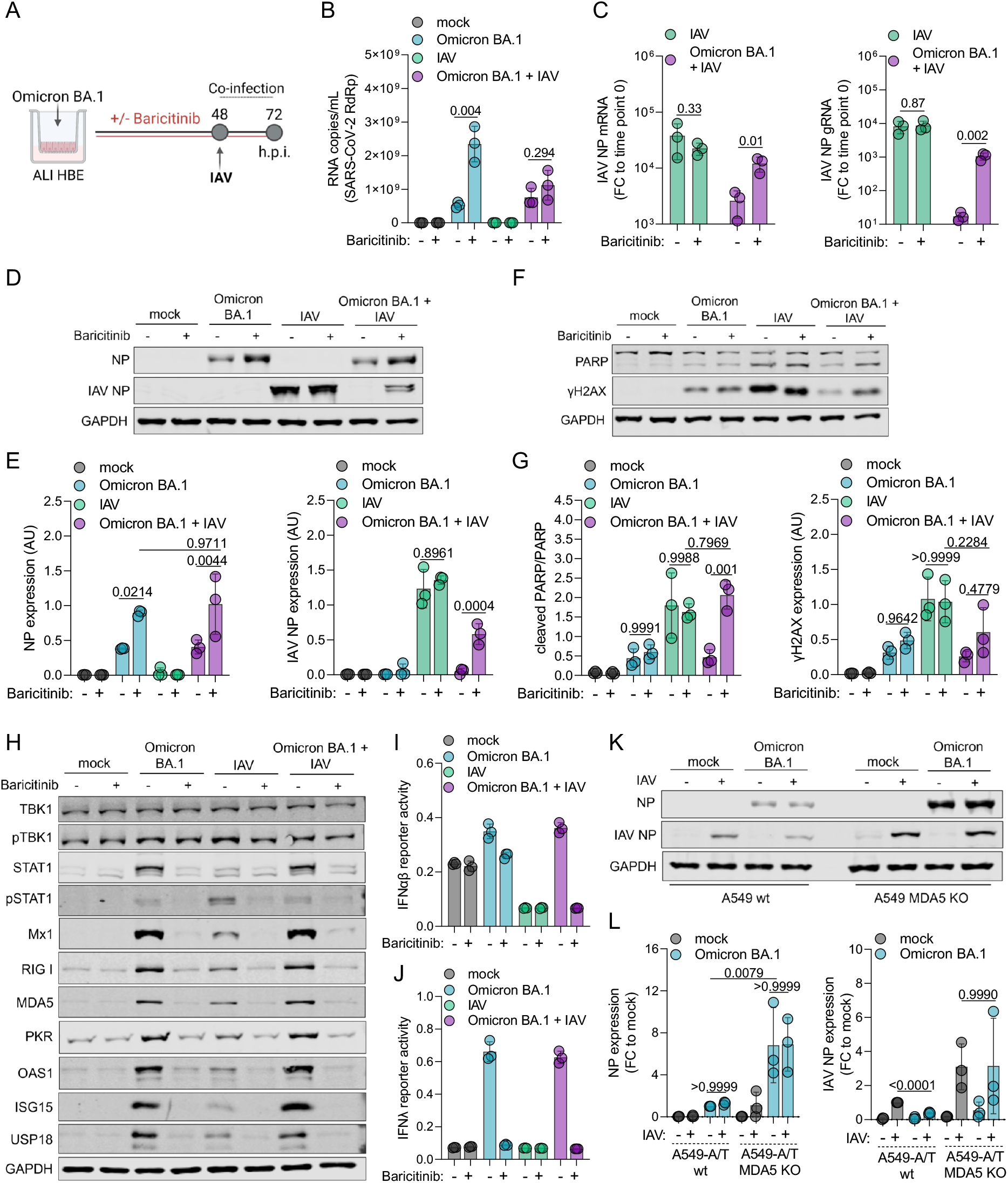
Inhibition of JAK/ STAT signalling prevents Omicron BA.1-mediated suppression of H1N1 influenza A virus (IAV) replication in air-liquid-interface (ALI) human bronchial epithelial (HBE) cultures. (A) Experimental set-up. (B) Genomic RNA copy numbers (RNA-dependent RNA polymerase gene = RdRp) in apical washes of BA.1 (MOI 1)-infected ALI HBE cultures 72h post infection in the absence or presence of baricitinib 1μM. Values represent mean ± SD of three biological replicates. P values were determined by Student’s t-test. (C) IAV NP mRNA (left) or genomic RNA (right) levels 24h post H1N1 (MOI 2) infection in the absence or presence of baricitinib 1μM. Bars represent mean ± SD of three biological replicates. P values were determined by Student’s t-test. (D) Immunoblot indicating BA.1 NP and IAV NP protein levels 72 h post BA.1 infection in the absence or presence of baricitinib 1μM. (E) Quantification of the immunoblot results from (D) by ImageJ. Values represent mean ± SD of three biological replicates. P values were calculated by two-way ANOVA. (F) Immunoblot indicating PARP cleavage and γH2AX protein levels 72 h post BA.1 infection in the absence or presence of baricitinib 1μM. (G) Quantification of the immunoblot results from (F) by ImageJ. Bars represent the quantification of the ratio between cleaved and total PARP (left) and cellular γH2AX levels (right). Values represent mean ± SD of three biological replicates. P values were calculated by two-way ANOVA. (H) Immunoblot displaying levels of proteins involved in interferon signalling in single- and co-infected ALI HBE cultures in the absence or presence of baricitinib 1μM. (I, J) Interferon (IFN)α/β (I) or -λ (J) activity in HEK reporter cell lines incubated with apical washes of ALI HBE cultures 72h post infection. (K) BA.1 NP and IAV NP protein levels in ACE2/ TMPRSS2-transduced A549 (A549-A/T) cells (A549-A/T wt) or A549-A/T MDA5 knock-out (KO) cells infected with BA.1 at MOI 0.01 for 24h and followed by influenza A virus (IAV) H1N1 (MOI 2) infection for an additional 24h. (L) Quantification of immunoblot results from (K) by ImageJ. Values represent the mean ± SD of three biological replicates. P values were calculated by two-way ANOVA.

In line with our previous findings, baricitinib also antagonised BA.1-induced suppression of H1N1-induced apoptosis as indicated by PARP cleavage and H1N1-induced DNA damage as indicated by cellular γH2AX levels (Figure 4F, Figure 4G). Furthermore, baricitinib abrogated the BA.1-mediated protection of the ALI HBE barrier integrity from H1N1-induced cytotoxicity (Suppl. Figure 2).

As indicated by the cellular levels of proteins involved in interferon signalling, the BA.1-induced interferon was not affected by of H1N1 (Figure 4H). Moreover, baricitinib inhibited interferon signalling in response to ALI HBE infection with either single virus and after co-infection with both viruses (Figure 4H) and suppressed interferon-α/β and -λ production (Figure 4I, Figure 4J).

The pattern recognition receptor MDA5 was previously shown to be critically involved in the SARS-CoV-2-mediated, and in particular the BA.1-mediated, interferon response [Yin et al., 2021; Bojkova et al., 2022; Bojkova et al., 2022a]. In agreement, BA.1-mediated inhibition of H1N1 infection was abrogated in MDA5 knock-out cells (Figure 4K, Figure 4L).

Taken together, our data show that BA.1-mediated suppression of H1N1 replication depends on the presence of MDA5 and is anatagonised by inhibition of JAK/ STAT signalling by baricitinib.

### Similar suppression of H1N1 influenza A virus replication by BA.1 and BA.5

Next, we compared the effects of BA.1 on interferon signalling and H1N1 replication to those of the Omicron subvariant BA.5 (Figure 5A), which is currently dominant in many parts of the world [Shrestha et al., 2022].

**Figure 5.**
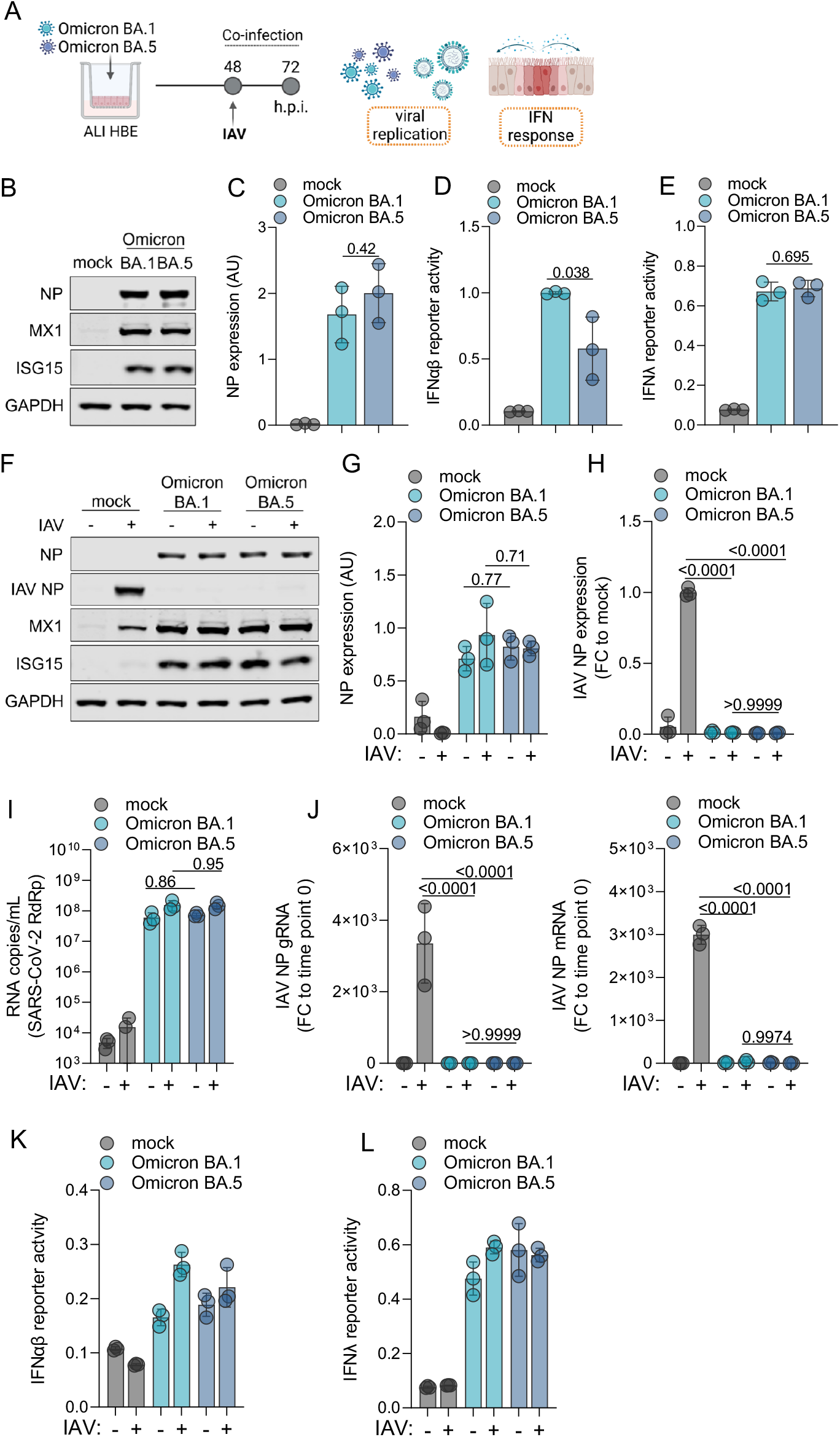
Effects of Omicron BA.1 and BA.5 on interferon signalling and H1N1 influenza A virus replication in air-liquid-interface (ALI) human bronchial epithelial (HBE) cell cultures. (A) Experimental design. (B) Cellular levels of the SARS-CoV-2 nucleoprotein (NP) and proteins involved in interferon signalling (MX1, ISG15) in BA.1- and BA.5 (MOI 1)-infected ALI HBE cultures 48h post infection. (C) Quantification of the NP immunoblot results from (B). Values represent mean ± SD from three biological replicates. P values were determined by Student’s t-test. (D,E) Interferon-α/β (D) or -λ (E) promotor activity in HEK reporter cell lines incubated with apical washes of BA.1- or BA.5-infected ALI HBE cultures 48h post infection. (F) SARS-CoV-2 NP (NP), influenza A virus NP (IAV NP), MX1, and ISG15 protein levels in BA.1 (MOI 1)-infected, BA.5 (MOI 1)-infected, IAV H1N1 (MOI 2)-infected, BA.1/ IAV co-infected, or BA.5/ IAV co-infected ALI HBE cultures. (G,H) Quantification of SARS-CoV-2 NP (G) and IAV NP (H) levels from (F) by ImageJ. (I) Genomic SARS-CoV-2 RNA (RNA-depended RNA polymerase/ RdRp gene) levels in BA.1-infected, BA.5-infected, BA.1/ IAV co-infected, and BA.5/ IAV co-infected cells 72h post infection. Values represent mean ± SD from three biological replicates. (J) Genomic IAV NP (gRNA) copy numbers and IAV NP mRNA levels in BA.1-infected, BA.5-infected, BA.1/ IAV co-infected, and BA.5/ IAV co-infected cells 72h post infection. (K, L) Interferon-α/β (K) or -λ (L) promotor activity in HEK reporter cell lines incubated with apical washes of BA.1-infected, BA.5-infected, BA.1/ IAV co-infected, and BA.5/ IAV co-infected ALI HBE cultures 72h post infection.

BA.1 and BA.5 infection of ALI HBE cultures resulted in similar SARS-CoV-2 NP protein levels (Figure 5B, Figure 5C) and induced similar interferon responses as indicated by the cellular levels of the interferon-stimulated gene products MX1 and ISG15 (Figure 5B) as well as interferon-α/β (Figure 5D) and -λ (Figure 5E) production in BA.1- and BA.5-infected ALI HBE cultures.

H1N1 co-infection did not significantly affect cellular SARS-CoV-2 NP levels or cellular MX1 and ISG15 levels (Figure 5F, Figure 5G). However, both Omicron subvariants suppressed H1N1 replication as indicated by cellular NP levels (Figure 5F, Figure 5H). These findings (limited impact of influenza A virus infection on SARS-CoV-2 replication, BA.1- and BA.5-mediated suppression of H1N1 replication) were confirmed by the determination of genomic SARS-CoV-2 RNA copy numbers (Figure 5I), genomic influenza A virus RNA copy numbers (Figure 5J), and H1N1 NP mRNA levels (Figure 5J). Both variants also induced similar interferon responses in the presence or absence of H1N1 as indicated by interferon-α/β (Figure 5K) and -λ (Figure 5L) production.

Taken together, BA.1 and BA.5 induce comparable interferon-mediated antiviral states in ALI HBE cultures that prevent H1N1 replication.

### BA.1-induced interferon signalling prevents H1N1 and H5N1 influenza A virus replication in primary human monocytes

Influenza A viruses can replicate in peripheral blood mononuclear cells, including CD14+ monocytes [Lersritwimanmaen et al., 2015; Lee et al., 2017], and interferon signalling in monocytes has been suggested to be a critical determinant of influenza A virus pathogenesis [Hartshorn, 2020; Fourati et al., 2022]. Highly pathogenic avian influenza A H5N1 virus has been described to replicate more readily in monocytes and macrophages than H1N1 [Yu et al., 2011; Hoeve et al., 2012; Cline et al., 2017; Lee et al., 2017; Lamichhane & Puthavathana, 2018; Lamichhane et al., 2018; Westenius et al., 2018]. Hence, we finally investigated the impact of BA.1 and Delta on H1N1 and H5N1 virus infection in primary human monocytes (Figure 6A).

**Figure 6.**
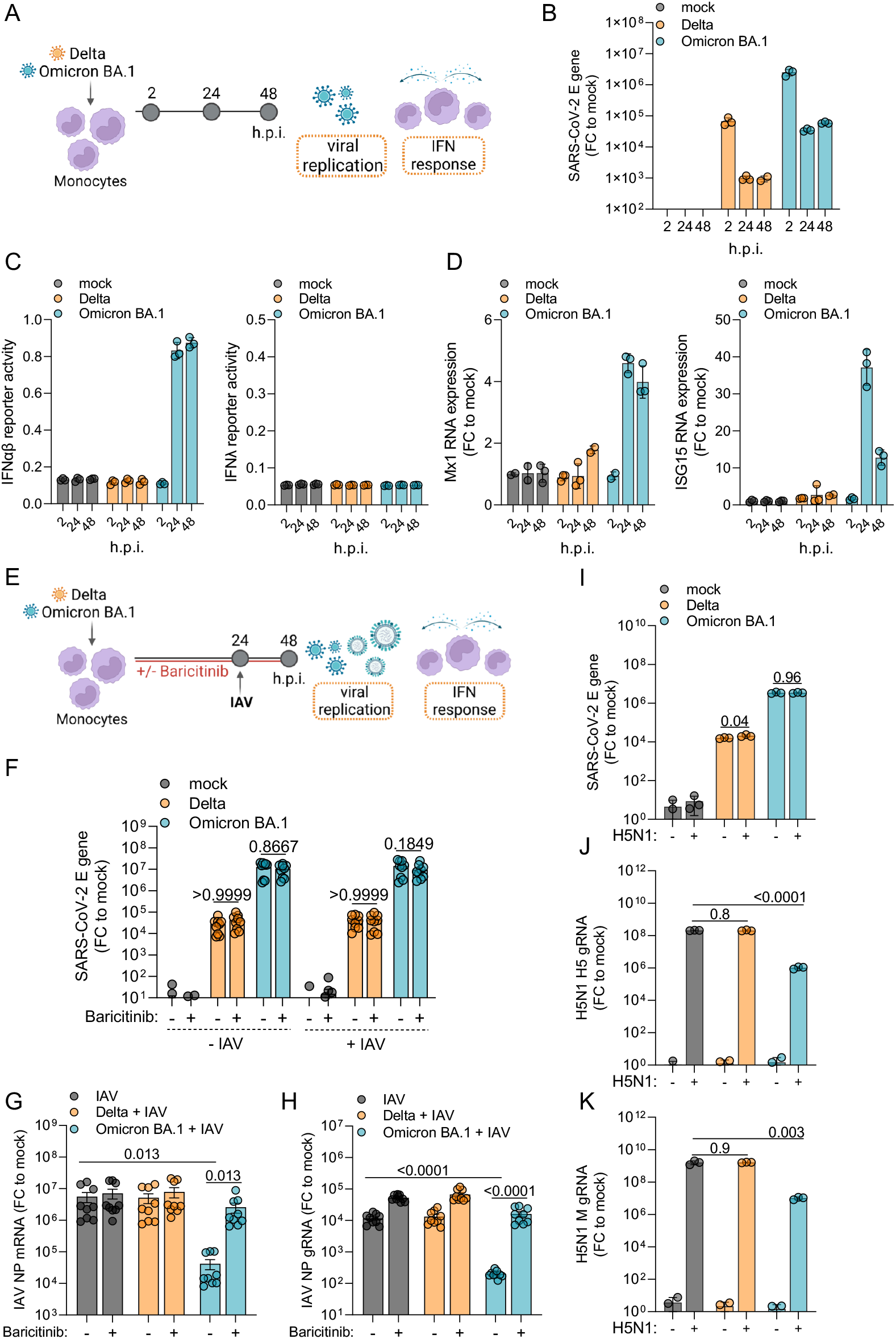
Impact of Omicron BA.1 and Delta on H1N1 and H5N1 influenza A virus (IAV) infection of primary human monocytes. (A) Experimental design underlying (B-D). (B) SARS-CoV-2 E gene expression levels as determined by quantitative PCR. Values represent mean ± SD of three biological replicates. (C) interferon-α/β (IFNα/β) (left) or IFNλ (right) activity induced by supernatants of Delta and BA.1 (MOI 1)-infected primary human monocytes in HEK-reporter cell lines. (D) Mx1 (left) and ISG15 (right) mRNA levels in Delta and BA.1 (MOI 1)-infected primary human monocytes as indicated by quantitative PCR. (E) Experimental design underlying (F-H). (F) SARS-CoV-2 E mRNA levels in monocytes infected with Delta (MOI 1), BA.1 (MOI 1), Delta plus H1N1 IAV (MOI 2), or BA.1 plus H1N1 IAV (MOI 2) in the absence or presence of baricitinib 1μM. P values were determined by Student’s t-test. (G, H) H1N1 IAV NP mRNA (G) or genomic mRNA (H) levels in monocytes infected with Delta (MOI 1), BA.1 (MOI 1), Delta plus H1N1 IAV (MOI 2), or BA.1 plus H1N1 IAV (MOI 2) in the absence or presence of baricitinib 1μM 24h post H1N1 IAV infection. P values were calculated by two-way ANOVA. (I-K) Impact of BA.1 and Delta infection on highly pathogenic avian H5N1 IAV virus infection in monocytes. SARS-CoV-2 genomic E RNA levels (I), genomic H5N1 H5 RNA levels (J), and genomic H5N1 IAV M levels (K) in monocytes infected with Delta (MOI 1), BA.1 (MOI 1), Delta plus H5N1 strain A/Vietnam/1203/04 (MOI 1), or BA.1 plus H5N1 (MOI 1). P values were calculated by two-way ANOVA. All values represent mean ± SD of monocyte preparations derived from three different donors (A-H) or one donor (I-K), which were each tested in three biological replicates.

The determination of SARS-CoV-2 E mRNA levels by quantitative PCR showed that BA.1 and Delta did not replicate in primary human monocytes (Figure 6B), which confirmed previous findings showing that SARS-CoV-2 causes abortive infections in monocytes [McLaughlin et al., 2021; Junqueira et al., 2022]. Only BA.1 induced a pronounced interferon response as indicated by interferon α/β signalling in a HEK reporter cell line incubated with supernatants from infected monocytes (Figure 6C), MX1 mRNA levels, and ISG15 mRNA levels (Figure 6D). In contrast to the findings in ALI HBE cultures (Figure 1H, Figure 2K, Figure 4J, Figure 5L), supernatants from BA.1-infected monocytes did not induce interferon-λ activity in a HEK reporter cell line (Figure 6C).

Next, we tested the impact of BA.1 and Delta on H1N1 replication in primary human monocytes in the absence or presence of the JAK inhibitor baricitinib (Figure 6E). Neither baricitinib nor H1N1 infection affect SARS-CoV-2 E mRNA levels (Figure 6F).

Similarly as in ALI HBE cultures, however, BA.1 (but not Delta) infection reduced H1N1 infection as indicated by H1N1 NP mRNA (Figure 6G) and H1N1 genomic NP RNA levels (Figure 6H), which was prevented by baricitinib (Figure 6G, Figure 6H).

Highly pathogenic H5N1 avian influenza A virus also did not affect SARS-CoV-2 levels in monocytes as indicated by SARS-CoV-2 genomic E RNA levels (Figure 6I). However, the determination of H5N1 virus genomic H5 (Figure 6J) and M (Figure 6K) levels showed that abortive BA.1 but not Delta infection inhibited H5N1 replication in primary human monocytes.

Taken together, abortive BA.1 but not Delta infection induced an interferon response and reduced H1N1 and H5N1 replication in monocytes. The JAK/ STAT inhibitor prevented BA.1-mediated H1N1 and H5N1 inhibition in monocytes, indicating that BA.1-mediated influenza A virus inhibition is caused by BA.1-induced interferon signalling.

### Comparison of the circulation of influenza-like illnesses during the Delta and BA.1 infection waves in England

An analysis comparing the spread of influenza-like illnesses (includes cases of clinically diagnosed influenza with or without a virus test confirmation) and COVID-19 in England showed that both Delta and influenza-like illnesses surged after all restrictions were removed on 19^th^ July 2021. When BA.1 became the dominant variant, however, the number of influenza-like illnesses strongly declined and has not surged since (Suppl. Figure 3). These data fit with our findings showing that BA.1 but not Delta induces an interferon response that prevents influenza A virus infection. Similarly, an analysis of data from the John Hopkins Microbiology Laboratory found a notable decrease in influenza A virus infections during the Omicron BA.1 wave between November 2021 and February 2022 [Eldesouki et al., 2022]. However, future research will have to analyse a possible causative relationship in more detail.

## Discussion

Here, we investigated the impact of Delta and the Omicron subvariants BA.1 and BA.5 on interferon signalling in ALI HBE cultures and primary human monocytes. Our results consistently showed that only Omicron causes pronounced production of biologically active type I and type III interferons (Suppl. Figure 4). The kinetics of the interferon type I (α/β) and type III (λ) responses provided additional mechanistic evidence explaining the reduced pathogenicity of Omicron relative to Delta [Wang et al., 2022]. BA.1 infection caused an early, transient interferon-α/β peak, which is anticipated to mediate a protective antiviral response but also to avoid deleterious inflammatory processes associated with prolonged interferon type I activation [Forero et al., 2019; King & Sprent, 2021]. Moreover, BA.1 infection caused a sustained interferon λ response known to inhibit virus replication and prevent excessive inflammation in the respiratory tract [Forero et al., 2019; Prokunina-Olsson et al., 2020; King & Sprent, 2021].

The Omicron-induced interferon response generated an antiviral state that protected infected cells from super-infection with influenza A viruses, showing that the Omicron-induced interferon response is of functional relevance. Our analysis of the spread of influenza-like illnesses during the Delta and BA.1 infection waves (Suppl. Figure 3) and data from the John Hopkins Microbiology Laboratory [Eldesouki et al., 2022] found that the levels of influenza A virus transmission levels strongly declined with the emergence of BA.1, but a causative relationship remains to be established.

Despite consistent BA.1-mediated interferon induction across the different cell types, the relative replication kinetics of Omicron and Delta differed between the models. BA.1 replicated less effectively than Delta in Caco-2F03 and Calu-3 [Bojkova et al., 2022; Bojkova et al., 2022a], but (in agreement with other findings [Hui et al., 2022]) faster than Delta in ALI HBE cultures. This probably reflects the contribution of many factors to SARS-CoV-2 replication. The spike (S) proteins of the BA.1 and Delta isolates that we used differ in 13 amino acid positions in the S receptor binding domain (Suppl. Figure 5). For example, BA.1 S is known to interact differently with its cellular receptor ACE2, to utilise ACE2 from a broader range of species as receptor, and to mediate increased virus uptake [Cameroni et al., 2022; Hong et al., 2022; Li et al., 2022; Meng et al., 2022; Rathnasinge et al., 2022; Willett et al., 2022; Zhang et al., 2022].

Increased BA.1 uptake may also contribute to the enhanced interferon response. Mechanistically, the BA.1-induced interferon response is, in agreement with previous studies in other cell types [Yin et al., 2021; Bojkova et al., 2022; Bojkova et al., 2022a], primarily mediated by the recognition of double-stranded RNA by the pattern recognition receptor MDA5, as suggested by the lack of a BA.1-induced interferon in MDA5 knock-out cells. In this context, virus uptake via the endosomal pathway was described to result in greater activation of pattern recognition receptors [Peacock et al., 2022].

BA.1 infection also induced an interferon-mediated antiviral state preventing influenza A (H1N1, H5N1) infection in monocytes despite only establishing an abortive infection, demonstrating that the antiviral state does not depend on a complete virus replication cycle resulting in the production of infectious virus. In agreement, UV-inactivated BA.1 had previously been shown to trigger a detectable interferon response in lung organ cultures, although this response was weaker than that induced by the replication-competent virus [Alfi et al., 2022].

The JAK/ STAT inhibitor baricitinib inhibited the Omicron-mediated antiviral response. This suggests that the Omicron-induced antiviral state is caused by MDA5-mediated production of interferon which activates interferon receptors that then trigger JAK/ STAT signalling [Li et al., 2020].

In addition to these findings, we also showed that BA.5, which is currently the dominant variant in many parts of the world [Shrestha et al., 2022], induces a comparable interferon response to BA.1 and that the BA.1- and BA.5-induced interferon responses protect infected cells from super-infection with influenza A viruses. The latter finding demonstrates that the BA.1- and BA.5-induced interferon signalling is of functional relevance. Baricitinib prevented the BA.1- and BA.5-induced suppression of influenza A virus replication, providing evidence that the BA.1- and BA.5-induced interferon response is responsible for the BA.1 and BA.5-induced influenza A virus inhibition.

In summary, we show that BA.1 and BA.5 (but not Delta) induce a functionally relevant pronounced interferon response that suppresses influenza A virus replication. Further research will have to show the relevance of our findings in the context of SARS-CoV-2/ influenza virus co-infection. Data on the severity of SARS-CoV-2/ influenza A virus co-infections are inconsistent in cell culture and animal models [Andrés et al., 2022; Kim EH et al., 2022; Kim HK et al., 2022; Oishi et al., 2022] and in humans [Cuadrado-Payán et al., 2020; Yue et al., 2020; Alosaimi et al., 2021; Stowe et al., 2021; Xiang et al., 2021; Krumbein et al., 2022; Swets et al., 2022]. This may not be a surprise, given the differences in interferon signalling between different SARS-CoV-2 variants that we present here and that were described in previous studies [Alfi et al., 2022; Bojkova et al., 2022; Bojkova et al., 2022a; Guo et al., 2022; Thorne et al., 2022]. Future studies may have to include more different virus variants and strains to establish a clearer picture.

In conclusion, our findings show that 1) BA.1 and BA.5 induce comparable interferon responses in ALI HBE cultures, 2) the Omicron-induced interferon-response is of functional relevance as it protects infected cells from influenza A virus replication, and 3) abortive BA.1 infection of monocytes is sufficient to produce a protective interferon response. Moreover, the kinetics of the Omicron-induced interferon response (early and transient type I response, sustained type III response) provide additional mechanistic evidence explaining why Omicron infections are usually associated with less severe disease than Delta infections.

## Methods

### Cell lines

HEK293 (HEK-Blue™ reporter cells, InvivoGen) -IFN-α/β and -IFN-λ cells were cultivated in DMEM (Gibco, ThermoFisher Scientific) with 10% heat-inactivated foetal bovine serum (Sigma), Pen-Strep (100 U/ml-100 μg/ml) (Sigma), 100 μg/ml Normocin™ (InvivoGen) and the required selective antibiotic for each cell line (IFN-α/β: Blasticidin and Zeocin; IFN-λ, Blasticidin, Puromycin, and Zeocin) at 37°C and 5% CO_2_.

A549-ACE2/TMPRSS2 cells (Invivogen) were grown in DMEM supplemented with 2 mM L-glutamine, 4.5 g/l glucose, 10% (v/v) heat-inactivated fetal bovine serum (FBS; 30 min at 56 °C), PenStrep (100 U/ml-100 μg/ml), 100 μg/ml Normocin, 10 μg/ml of Blasticidin, 10 μg/ml of Blasticidin, 100 μg/ml of Hygromycin, 0.5 μg/ml of Puromycin, and 100 μg/ml of Zeocin. A549-ACE2/TMPRSS2 MDA5 KO cells (Invivogen) were grown in DMEM supplemented with 2 mM L-glutamine, 4.5 g/l glucose, 10% (v/v) heat-inactivated fetal bovine serum (FBS; 30 min at 56 °C), PenStrep (100 U/ml-100 μg/ml), 100 μg/ml Normocin, 10 μg/ml of Blasticidin, 100 μg/ml of Hygromycin, 0.5 μg/ml of Puromycin, and 100 μg/ml of Zeocin.

All cell lines were regularly tested for mycoplasma contamination.

### Air-liquid interface cultures

Lung tissue for the isolation of primary epithelial cells was provided by the Hannover Medical School, Institute of Pathology (Hannover, Germany). The use of tissue was approved by the ethics committee of the Hannover Medical School (MHH, Hannover, Germany, number 2701–2015) and was in compliance with The Code of Ethics of the World Medical Association. Primary bronchial epithelial cells were isolated from the lung explant tissue of a patient with lung emphysema as described previously [van Wetering et al., 2000]. All patients or their next of kin gave written informed consent for the use of their lung tissue for research.

Basal cells were expanded in Keratinocyte-SFM medium supplemented with bovine pituitary extract (25 μg/mL), human recombinant epidermal growth factor (0.2 ng/mL, all from Gibco, Schwerte, Germany), isoproterenol (1 nM, Sigma), Antibiotic/Antimycotic Solution (Sigma-Aldrich), and MycoZap Plus PR (Lonza, Cologne, Germany) and cryopreserved until further use.

For differentiation, the cells were thawed and passaged once in PneumaCult-Ex Medium (StemCell Technologies, Cologne, Germany) and then seeded on transwell inserts (12-well plate, Sarstedt, Nümbrecht, Germany) at 4 × 10^4^ cells/insert. Once the cell layers reached confluency, the medium on the apical side of the transwell was removed, and medium in the basal chamber was replaced with PneumaCult ALI Maintenance Medium (StemCell Technologies), including Antibiotic/Antimycotic Solution (Sigma-Aldrich) and MycoZap Plus PR (Lonza). During a period of four weeks, the medium was changed and the cell layers were washed with PBS every other day. Criteria for successful differentiation were the development of ciliated cells and ciliary movement, an increase in transepithelial electric resistance indicative of the formation of tight junctions, and mucus production.

### SARS-CoV-2 variants preparation

The SARS-CoV-2 isolates Omicron BA.1 (B.1.1.529: FFM-SIM0550/2021, EPI_ISL_6959871, GenBank ID OL800702), Delta (B.1.167.2: FFM-IND8424/2021, GenBank ID MZ315141), and Omicron BA.5 (GenBank ID OP062267) were isolated in Caco-2-F03 cells as previously described [Cinatl et al., 2004; Bojkova et al., 2021] and stored at −80°C. All variants underwent maximum two passages. Virus titres were determined as TCID50/mL.

### Influenza A virus strains H1N1 and H5N1

The H1N1 influenza strain A/NewCaledonia/20/99 was received from the World Health organisation (WHO) Influenza Centre (National Institute for Medical Research, London, UK) and stocks were prepared by cultivation on MDCK cells (ATCC, CCL-34) in medium containing 2μg/ml trypsine. Virus stocks were stored at −80 °C. Virus titres were determined as TCID50/mL.

The H5N1 influenza strain A/Vietnam/1203/04 was received from the World Health organisation (WHO) Influenza Centre (National Institute for Medical Research, London, UK). Virus stocks were prepared by infecting Vero cells, and aliquots were stored at −80 °C. Virus titres were determined as TCID50/mL.

### Barrier integrity measurement

For trans-epithelial electrical resistance (TEER) measurement, medium was added to the apical side 30min prior to measurement with a chopstick electrode connected to a Volt-Ohm-meter (Millicell^®^ ERS-2, Merck, Darmstadt, Germany) according to the manufacturer’s instructions. Blank inserts served as baseline. The apical medium was removed after the measurement.

### Activation of caspase 3/7

Caspase 3/7 activity was measured using the Caspase-Glo assay kit (Promega, Madison, WI, USA), according to the manufacturer’s instructions. Briefly, 100 μL of Caspase-Glo reagent were added to each well containing 100 μL of tested sample, mixed, and incubated at room temperature for 30 min. Luminescence intensity was measured using an Infinite M200 microplate reader (Tecan).

### Co-infection assay in ALI HBE

ALI HBE cultures were infected with SARS-CoV-2 variant at MOI 1 from the apical site. The inoculum was incubated for 2 h, then removed and cells were washed three times with PBS. H1N1 A/NewCaledonia/20/99 at MOI 2 was added 48 h post SARS-CoV-2 infection. The inoculum was incubated for 2 h, then removed and cells were washed three times with PBS.

### Detection of extracellular and intracellular RNA

SARS-CoV-2 RNA from the apical washes of the ALI HBE culture was isolated using QIAamp Viral RNA Kit (Qiagen, Hilden, Germany) according to the manufacturer’s instructions. RNA was subjected to OneStep qRT-PCR analysis using the Luna Universal One-Step RT-qPCR Kit (New England Biolabs, Frankfurt am Main, Germany) and a CFX96 Real-Time System, C1000 Touch Thermal Cycler (Bio-Rad, Feldkirchen, Germany). Primers were adapted from the WHO protocol29 targeting the open reading frame for RNA-dependent RNA polymerase (RdRp): RdRP_SARSr-F2 (GTG ARA TGG TCA TGT GTG GCG G) and RdRP_SARSr-R1 (CAR ATG TTA AAS ACA CTA TTA GCA TA) using 0.4 μM per reaction. Standard curves were created using plasmid DNA (pEX-A128-RdRP) as previously described [Bojkova et al., 2020].

Intracellular RNA isolation was carried out according to manufacturer’s protocol (RNeasy 96 QIAcube HT Kit, Qiagen, Hilden, Germany). Detection of selected targets was performed with Luna^®^ Universal One-Step RT-qPCR (New England BioLabs Inc.) according to manufacturers protocol using specific primers: TBP (fw:5’-ATCAGAACAACAGCCTGCC-3’; rev: 5’-GGTCAGTCCAGTGCCATAAG-3’); SARS-CoV-2 E gene (fw:5’-ACAGGTACGTTAATAGTTAATAGCGT-3’; rev:5’-ATATTGCAGCAGTACGCACACA-3’); ISG15 (fw:5’-GAGAGGCAGCGAACTCATCT-3’; rev:5’-AGGGAC ACCTGGAATTCGTT-3’); MX1 (fw:5’-TTTTCAAGAAGGAGGCCAGCAA-3’; rev:5’-TCAGGAACTTCCGCTTGTCG-3’); H5N1 H5 gene (fw:5’-GCCATTCCACAACATACACCC-3’; rev:5’-CTCCCCTGCTCATTGCTATG-3’); H5N1 M gene (fw:5’-TTCTAACCGAGGTCGAAACG-3’; rev:5’-ACAAAGCGTCTACGCTGCAG-3’); IAV NP-mRNA (fw:5’-GACTCACATGATGATCTGGCA-3’; rev:5’-CTTGTTCTCCGTCCATTCTCA-3’); IAV NP-gRNA (fw:5’-AACGGCTGGTCTGACTCACATGAT-3’; rev:5’-AGTGAGCACATCCTGGGATCCATT-3’).

### Immunoblot analysis

Whole-cell lysates were prepared using Triton-X sample buffer containing protease inhibitor cocktail (Roche). The protein concentration was assessed by using DC Protein assay reagent (Bio-Rad Laboratories). Equal protein loads were separated by sodium dodecyl sulfate-polyacrylamide gel electrophoresis and proteins were transferred to nitrocellulose membranes (Thermo Scientific). For protein detection the following primary antibodies were used at the indicated dilutions: GAPDH (Cell Signaling, #2118, 1:4000), γH2AX (Cell Signaling, #9718, 1:1000), H1N1 (Influenza A Virus) Nucleoprotein (Bioss, #bs-4976R, 1:4000), ISG15 (Santa Cruz Biotechnology, #sc-166755, 1:200), MDA5 (Cell Signaling, #5321, 1:1000), Mx1 (Cell Signaling, #37849, 1:1000), OAS1 (Cell Signaling, #14498, 1:1000), PARP (Cell Signaling, #9542, 1:1000), PKR (Cell Signaling, #12297, 1:1000), SARS-CoV-2 Nucleocapsid (Sino Biological, #40143-R019, 1:10000), STAT1 (Cell Signaling, #9172, 1:1000), phospho-STAT1 Y701 (Cell Signaling, #9171, 1:1000), TBK1 (Cell Signaling, #3013, 1:1000), phospho-TBK1 S172 (Cell Signaling, #5483, 1:1000), USP18 (Cell Signaling, #4813, 1:2000) and RIG1 (Cell Signaling, #3743, 1:1000). Protein bands were visualised using IRDye-labeled secondary antibodies at dilution 1:40000 (LI-COR Biotechnology, IRDye^®^800CW Goat anti-Rabbit, #926-32211 and IRDye^®^800CW Goat anti-Mouse IgG, #926-32210) and Odyssey Infrared Imaging System (LI-COR Biosciences).

### Detection of type I and type III IFNs production

Detection of type I and type III IFNs in supernatants was carried out with HEK-Blue™ IFN-α/β (type I) and HEK-Blue™ IFN-λ (type III) cells according to manufacturer’s protocol. HEK cells were washed twice with PBS and detached in presence of PBS by tapping the flask. Cells were subsequently centrifuged at 200g for 5 min and resuspended in Test Medium (DMEM, 4.5 g/l glucose, 2 mM L-glutamine, 10% (v/v) heat-inactivated FBS, 50 U/ml penicillin, 50 μg/ml streptomycin, 100 μg/ml Normocin™) at 280.000 cells/ml. 20μl supernatant were added in 96 well plates and 180 μl cell suspension was added to the wells. Cells and supernatant were incubated at 37°C and 5% CO_2_ for 24h. After incubation, 20 μl supernatant was removed and incubated with 180 μl QUANTI-Blue™ Solution for 1-3 h. SEAP (secreted embryonic alkaline phosphatase) levels were determined using a spectrophotometer at 620nm.

### PBMCS isolation

Human peripheral blood mononuclear cells were isolated from buffy coats of healthy donors (RK-Blutspendedienst Baden-Württemberg-Hessen, Institut für Transfusionsmedizin und Immunhämatologie Frankfurt am Main, Germany). After centrifugation on a Ficoll (Pancoll, PAN-Biotech, Aidenbach, Germany) density gradient, mononuclear cells were collected from the interface, washed with PBS, and plated on cell culture dishes (Cell+, Saarstedt, Nümbrecht, Germany) in RPMI1640 (Gibco, ThermoFisher Scientific, Waltham, MA, USA) supplemented with 100 IU/mL penicillin and 100 g/mL streptomycin. After incubation for 90 min(37°C, 5% CO2), non-adherent cells were removed, and the medium was changed to RPMI1640 supplemented with 100 IU/mL penicillin, 100 μg/mL of streptomycin, and 3% human serum (RK-Blutspendedienst Baden-Württemberg-Hessen, Institut für Transfusionsmedizin und Immunhämatologie Frankfurt am Main, Germany).

### Co-infection assay in human PBMCs

Human PBMCs were infected with SARS-CoV-2 variants at MOI of 1 for 2 h in infection medium (RPMI1640 supplemented with 100 IU/mL penicillin and 100 g/mL streptomycin, 1% heat-inactivated fetal bovine serum) at 37°C at 5% CO2. Afterwards the cells were washed twice with PBS and incubated for 24 h in infection medium. For co-infection, the cells were either infected with influenza A virus H1N1/New Caledonia/20/99 (MOI 2), influenza A virus H5N1 A/Vietnam/1203/04 (MOI 1) or treated with medium for 2 h, washed twice and incubated again for 24 h. After each washing step and after 48 h supernatant samples were taken and cells were lysed for RNA isolation.

### Statistics

Results are expressed as the mean ± standard deviation (SD) of the number of biological replicates indicated in figure legends. Statistical significance is depicted directly in graphs and the statistical tests used for the calculation of p values are indicated in the figure legends. GraphPad Prism 9 was used for visualisation of the data and for calculation of statistical significance.

## Acknowledgements

We thank Lena Stegman, Kerstin Euler, and Sebastian Grothe for their technical assistance.

