## Supplementary material for "Omicron-induced interferon signalling prevents influenza A virus infection": Suppl. Figure 1

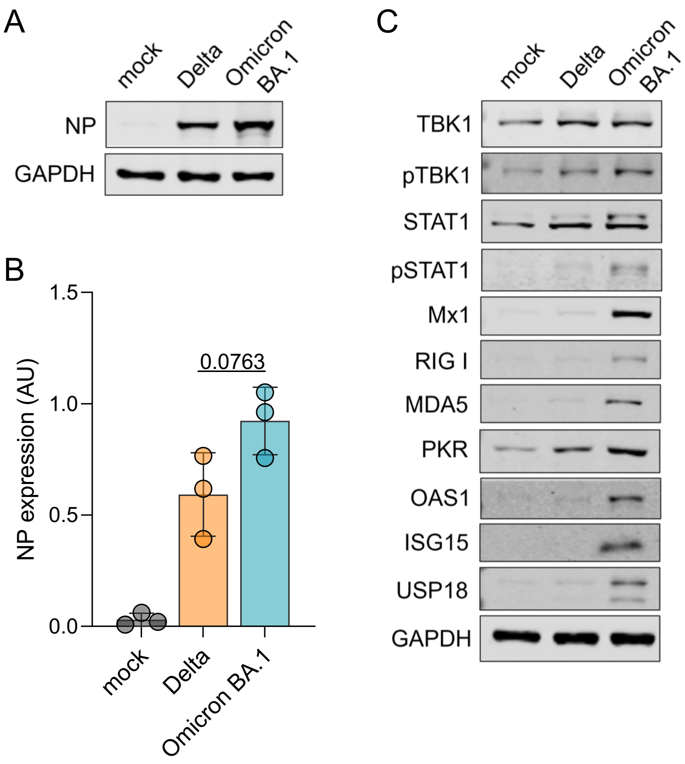


**Supplementary Figure 1. Infection rate and interferon response in Delta- and BA.1-infected air-liquid-interface (ALI) human bronchial epithelial (HBE) cell cultures prior to influenza A virus infection.** (A) Immunoblot of SARS-CoV-2 NP expression levels in ALI HBE cultures 48h post infection (MOI 1). (B) Quantification NP immunoblots. Bars display mean ± SD of three biological replicates. P values were calculated by Student´s t-test. (C) Immunoblot of proteins involved in the interferon response in ALI HBE cultures 48 post infection with BA.1 or Delta (MOI 1).
