## Supplementary material for "Omicron-induced interferon signalling prevents influenza A virus infection": Suppl. Figure 2

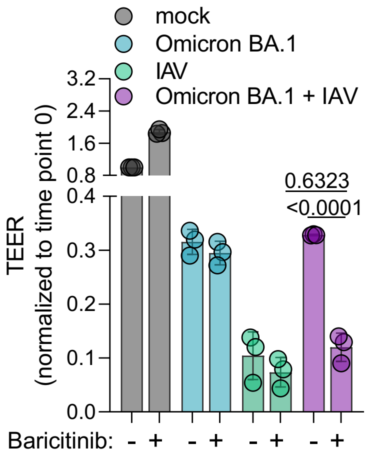


**Supplementary Figure 2. Impact of BA.1 and influenza A virus (IAV) infection on the barrier integrity of air-liquid-interface (ALI) human bronchial epithelial (HBE) cell cultures in the presence or absence of baricitinib.** Cells were infected as described in Figure 4 and the transepithelial electric resistance (TEER) was measured 72h post infection with BA.1. Values represent mean ± SD of three biological replicates. P values were calculated by two-way ANOVA.
