## Supplementary material for "Omicron-induced interferon signalling prevents influenza A virus infection": Suppl. Figure 3

A

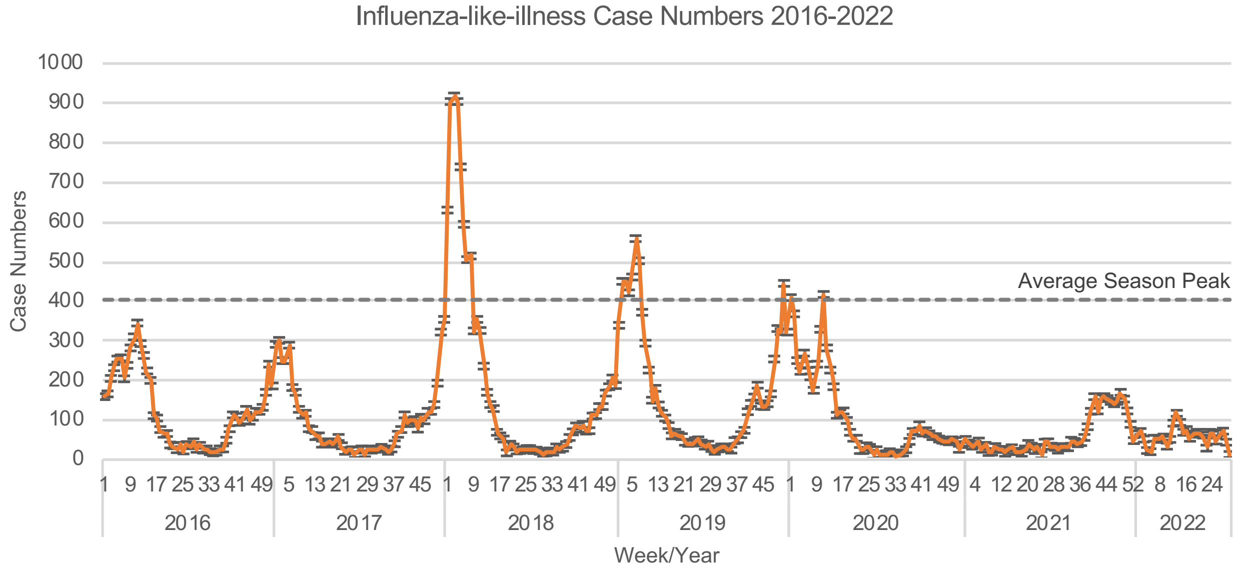


B

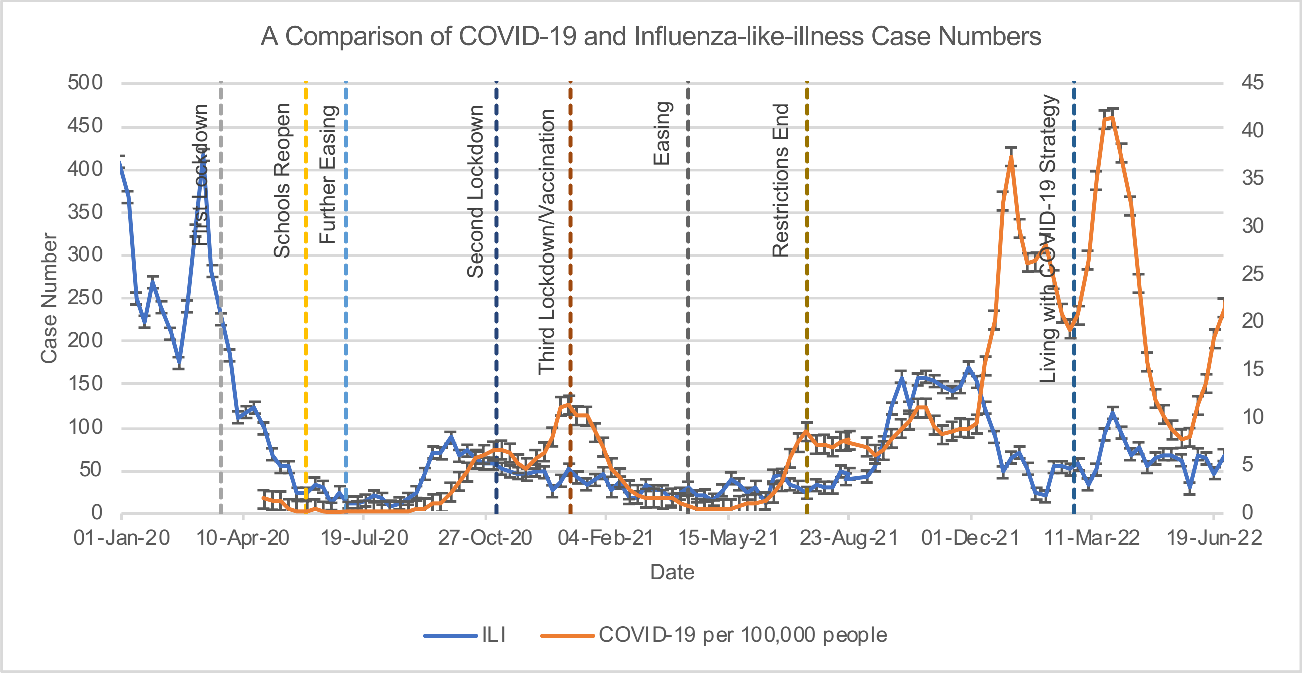

Suppl. Figure 3

C

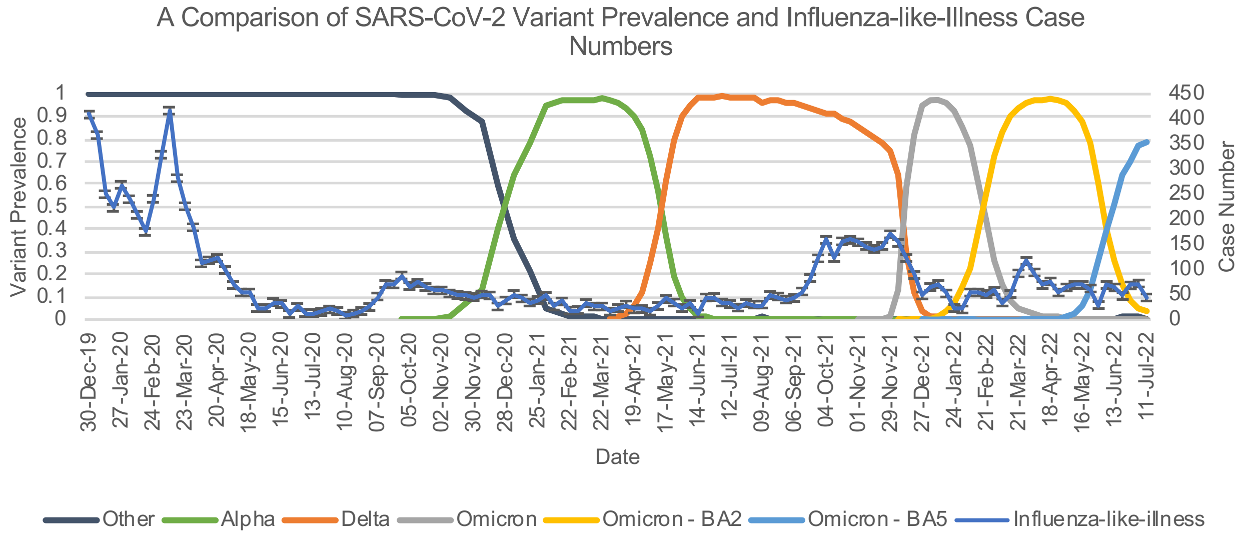


**Supplementary Figure 3. Cases of influenza-like illnesses and COVID-19 in England.** A) Influenza-like illness (clinically diagnosed influenza cases with or without a virus test confirmation) weekly case numbers plotted against week and year in which they occurred. Standard errors for each are week shown. The average of the highest case numbers of Influenza-like illness from the 2016-2020 seasons is indicated with a dashed line.

B) Comparison of COVID-19 and Influenza-like illness case numbers. Influenza-like illness case numbers (blue) refer to the primary (left) y axis. Estimated average number of positive COVID-19 cases per 100,000 people (orange) refer to the secondary (right) y axis. Weekly case numbers plotted against week in which they occurred, with dates spanning 30th December 2019 to 18th July 2022. Standard errors are shown for both datasets. COVID-19 prevention measures are indicated with dashed lines at the dates when they were imposed or removed.

C) SARS-CoV-2 variant prevalence and Influenza-like illness case numbers in England. SARS-CoV-2 variant prevalence refers to the primary (left) y axis and Influenza-like illness case numbers to the secondary (right) y axis. Standard errors are presented for Influenza-like illness case numbers.

Influenza-like illness case data are derived from Royal Collage of General Practitioners (RCGP) Research and Surveillance Centre (RSC) public health data (https://www.rcgp.org.uk/representing-you/research-at-rcgp/research-surveillance-centre/public-health-data, accessed on 16th August 2022). COVID-19 case numbers were derived from the Coronavirus (COVID-19) Infection Survey: England - Office for National Statistics (https://www.ons.gov.uk/peoplepopulationandcommunity/healthandsocialcare/conditionsanddiseases/datasets/coronaviruscovid19infectionsurveydata, accessed on 17th August 2022). Prevention measure date and description aided by Aspinall E. COVID-19 Timeline. British Foreign Policy Group. 2020 (https://bfpg.co.uk/2020/04/covid-19-timeline/, accessed on 17th August 2022). SARS-CoV-2 variant prevalence was extracted from Investigation of SARS-CoV-2 variants: technical briefings, GOV.UK (https://www.gov.uk/government/publications/investigation-of-sars-cov-2-variants-technical-briefings, accessed on 16th August 2022).
