## Supplementary material for "Omicron-induced interferon signalling prevents influenza A virus infection": Suppl. Figure 4

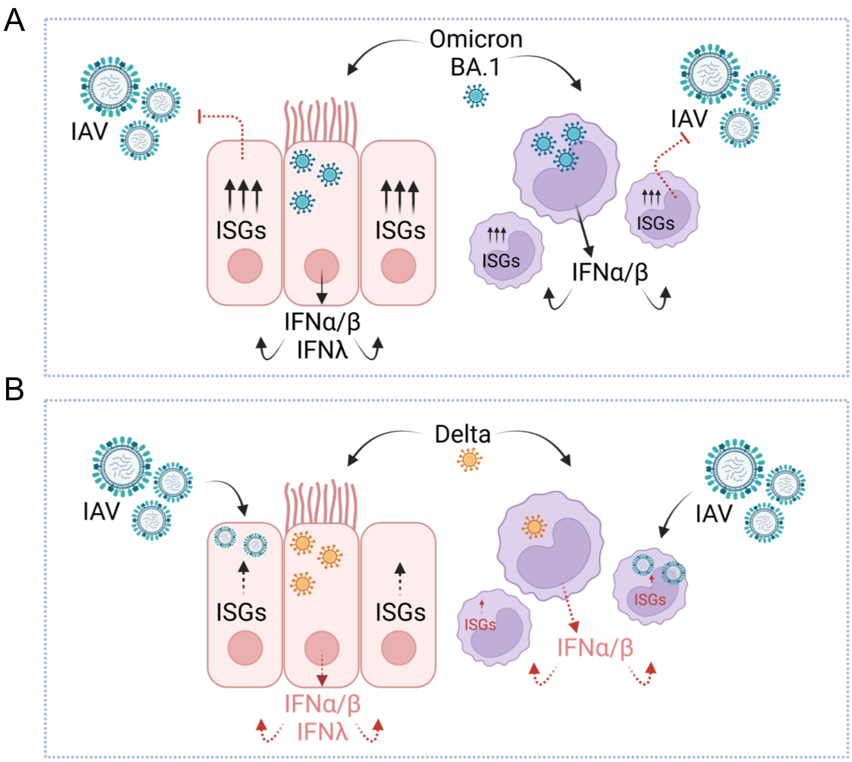


**Supplementary Figure 4. Overview of the study findings.** SARS-CoV-2 Omicron BA.1 and BA.5 (A) but not Delta (B) induce an interferon-mediated antiviral state that protects infected cells from influenza A virus infection.
