## Supplementary material for "Omicron-induced interferon signalling prevents influenza A virus infection": Suppl. Figure 5

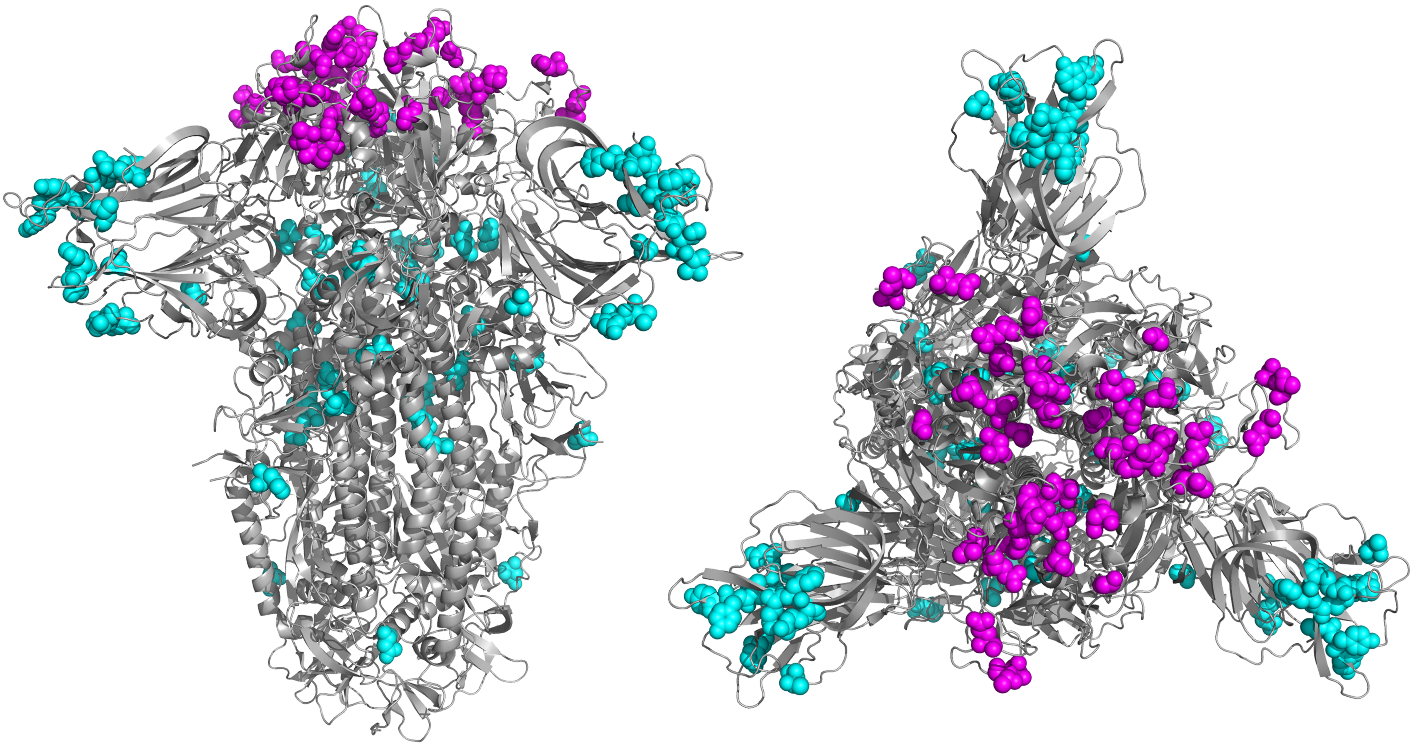

**Supplementary Figure 5. Amino acid changes in the spike (S) protein between the Omicron BA.1 and the Delta isolate used in this study.** The S protein is shown from two different angles. Sequence variants located in the receptor binding domain are highlighted in magenta. Changes to residue bonding were visualised using Pymol (https://pymol.org/2/) using Protein Databank in Europe (PDBe) structure 7fg3.
